# Detection of statistically robust interactions from diverse RNA-DNA ligation data

**DOI:** 10.1101/2024.09.17.610461

**Authors:** Simonida Zehr, Sandra Seredinski, Emma C. Walsh, Alessandro Bonetti, Matthias S. Leisegang, Ralf P. Brandes, Marcel H. Schulz, Timothy Warwick

## Abstract

Chromatin-localized RNAs play diverse roles in gene regulation and nuclear architecture. Mapping genome-wide RNA-DNA interactions is possible using a variety of molecular methods, including using bridging oligonucleotides to ligate RNA and DNA in proximity. While molecular methods have progressed, a robust computational method for calling biologically meaningful RNA-DNA interactions from these data is lacking. Herein, we present *RADIAnT*, a reads-to-interactions pipeline for analyzing RNA-DNA ligation data. *RADIAnT* calls interactions against a dataset-specific, unified background which considers RNA binding site-TSS distance and genomic region bias. By scaling the background by RNA abundance, *RADIAnT* is sensitive enough to detect specific interactions of lowly expressed transcripts, while remaining specific enough to discount false positive interactions of highly abundant RNAs. *RADIAnT* outperforms previously proposed methods in the accurate recall of genome-wide *Malat1*-DNA interactions, and in a use case, was utilized to identify dynamic chromatin-associated RNAs in the physiologically- and pathologically-relevant process of endothelial-to-mesenchymal transition.

## Main

The recognition of RNA as a gene regulatory molecule has prompted research into uncovering the exact mechanisms involved [1]. In some cases, it has been demonstrated that RNAs associate with chromatin in order to exert their regulatory effects [2]. Consequently, several molecular methods have been developed to identify the DNA binding sites of either one or multiple RNAs. These include, but are not limited to, antisense oligonucleotide-based approaches (ChIRP-seq [3], ChOP-seq [4], CHART-seq [5], RAP-DNA [6]), immunoprecipitation-based approaches (triplexDNA-seq [7]), and more recently oligonucleotide linker-based methods.

Linker-based methodologies for detection of all-to-all RNA-DNA interactions rely on the addition of labelled bridging oligonucleotides to disrupted nuclei (**Fig. 1a**). Assays belonging to this class of methods include RADICL-seq [8], Red-C [9], GRID-seq [10], ChAR-seq [11] and iMARGI [12]. While technical details vary, the end results of these methods are generally chimeric RNA-DNA sequencing reads separated by a bridging oligonucleotide, which are then subjected to library preparation and sequencing. Resulting reads can be computationally decomposed into paired RNA and DNA sequences. Once aligned to the genome, the mapped RNA and DNA permit identification of RNA molecules and genomic loci that were in proximity at the time of the experiment.

**Fig. 1.**
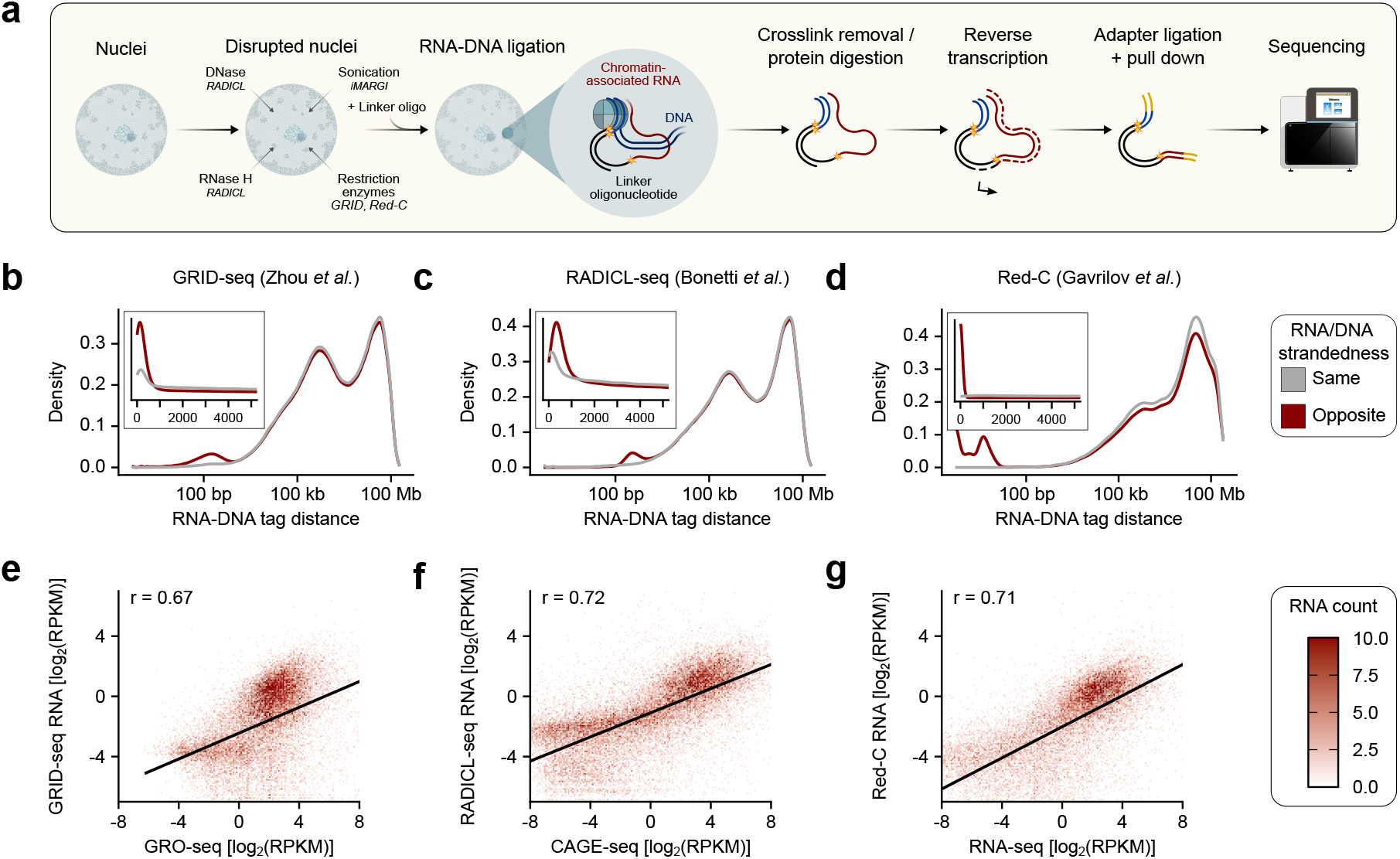
RNA-DNA ligation data are confounded by distance and RNA abundance. **(a)** Generalized workflow for RNA-DNA ligation experiments. **(b-d)** RNA-DNA tag distances in base pairs from published GRID-seq [18], RADICL-seq [8] and Red-C [9] experiments, split by stranded-ness. **(e-g)** Correlation between RNA abundance in RNA-DNA ligation data and RNA expression as captured by GRO-seq, CAGE or RNA-seq.

RNA-DNA ligation data have the potential to be highly complex. The theoretical search space which these data cover is between all RNA molecules in the cell, in combination with the entire length of the genome. In this sense, one would expect the data to be similar to Hi-C [13], the all-to-all method for capturing DNA-DNA interactions. However, in Hi-C, the linear distance between two interacting regions of the genome can always be calculated. These distances are often used in the building of statistical backgrounds, against which overrepresented interactions can be called [14]. Given the theoretical capability of RNA molecules to diffuse across nuclear space [15], building an appropriate statistical background is a more difficult prospect in RNA-DNA ligation data.

Despite this, several methods for the computational analysis of RNA-DNA ligation data have been proposed. The RADICL-seq publication [8] suggests calling RNA-DNA interactions using a binomial test, comparing the mean RNA interaction count at a given genomic bin compared to the observed count. In GRID-seq [10], the authors propose building a background based on *trans*-chromsomal mRNA-DNA interaction frequencies, against which the fold enrichment of RNA-DNA interactions is calculated. This approach is also suggested by the authors of Red-C [9]. More recently, the peak-calling approach *BaRDIC* was published [16], which considers gene-interaction distance and variation in RNA-binding profiles to detect genome-wide RNA-DNA interactions from all-to-all RNA-DNA ligation data. Similarly to GRID-seq, *BaRDIC* utilizes mRNA-DNA contacts to build a statistical background. For interactions in *cis*, this background is additionally scaled by the distance between the RNA transcription locus and interacting region of the genome. At present, there exists no standardized analysis for RNA-DNA ligation data that can effectively manage confounding factors such as distance and abundance and eliminate experimental noise, while still maintaining sufficient sensitivity to detect biologically meaningful RNA-DNA interactions.

Here, we present *RADIAnT* (RNA And DNA Interactions Analyzed ‘n’ Tested), a statistical method for calling robust interactions from diverse RNA-DNA ligation data types. By considering and accounting for the main confounding factors in these data, *RADIAnT* is capable of detecting known RNA-DNA interactions with high specificity, whilst simultaneously remaining sensitive enough to identify interactions of lowly-expressed RNAs. *RADIAnT* detects specific RNA-DNA interactions across different input data consistently, and outperforms other computational methods. This is exemplified by the accurate recall of *Malat1*-DNA interactions previously reported in one-to-all data [17]. When implemented on newly-generated RADICL-seq data from endothelial-to-mesenchymal transition, *RADIAnT* was used to identify putative process-specific RNA regulators, whose modulation has the potential to affect this disease-relevant process. *RADIAnT* is incorporated into a complete reads-to-interactions pipeline, which can be implemented on any RNA-DNA ligation data with minimal pre-processing requirements.

## Results

### RNA-DNA ligation data are confounded by distance and abundance

Published RADICL-seq [8], GRID-seq [10] and Red-C [9] datasets were downloaded, split into paired RNA and DNA sequences, and aligned to appropriate genome assemblies. For each RNA-DNA read pair which mapped to the same chromosome, the linear distance between the mapped RNA and DNA was calculated in base pairs. Upon plotting the density of RNA-DNA distances by strandedness, a clear enrichment could be observed in RNA-DNA distances smaller than 1 kilobase where strandedness was opposite (**Fig. 1b-d**). This corresponds to a high proportion of RNA associating with complementary DNA in its own transcriptional locus, which is likely a result of nascent transcription.

Given the potential impact of active transcription on the data, the relationship between RNA expression and abundance in RNA-DNA ligation data was also investigated. For each ligation method, accompanying gene expression data was also downloaded and analyzed. In each case, significant positive correlations between RNA expression and abundance in the ligation data could be observed (**Fig. 1e-g**).

These two observations made it clear that both the abundance of the RNA and the RNA gene locus-binding site distance should be considered when analyzing these data. Additionally, to better control for variation between methods and potential quality differences between experiments, the statistical background should be built from the given dataset, rather than in a generalized manner.

### *RADIAnT* calls RNA-DNA interactions against a unified, dataset-specific background

Based on the observations described above, we formulated *RADIAnT. RADIAnT* uses the RNA-DNA ligation data provided by the user to build a unified, dataset-specific statistical background, against which RNA-DNA interactions are called. To aggregate interaction counts and enable quantification of the data, the genome is split into bins of a user-defined size (e.g. 5 kb). RNA-DNA interaction counts are then summarized as the total reads between an RNA and a genomic bin. By dividing the RNA-bin interaction count by the total counts for the RNA in question, the interaction frequency of the RNA at the bin can be derived, a metric which is robust to expression differences between RNAs.

To account for the effect of nascent transcription, a distance-specific background is constructed from the input dataset. This is done by calculating the mean RNA-bin interaction frequency at defined linear distances from the RNA gene locus (**Fig. 2a, top**). The mean of these frequencies per distance is then taken as the distance-specific *RADIAnT* background for the experiment in question.

**Fig. 2.**
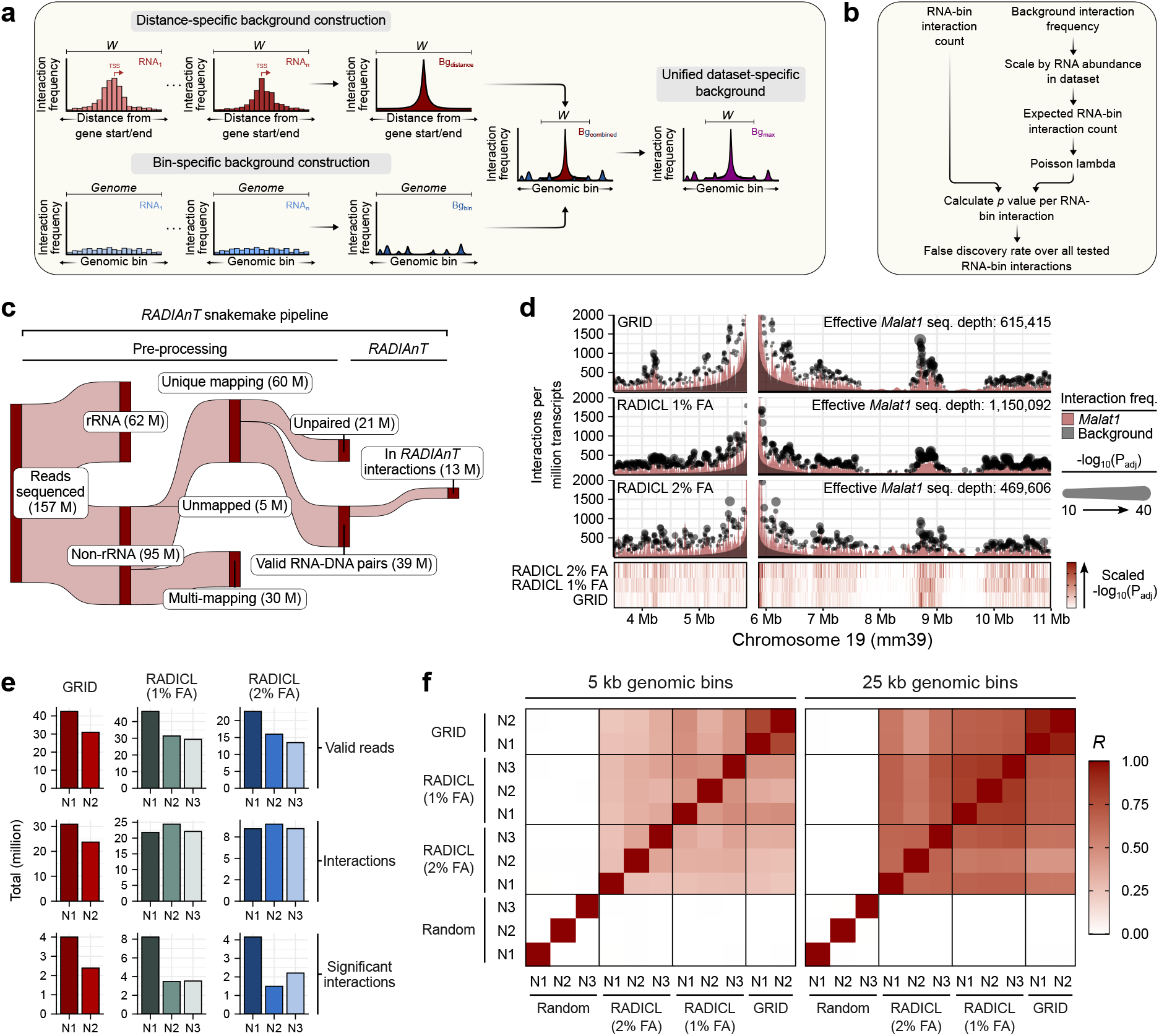
*RADIAnT* calls consistent RNA-DNA interactions regardless of input data type. **(a-b)** *RADIAnT* utilised the input data to build distance- and bin-specific statistical backgrounds, which are merged into a single unified background against which RNA-DNA interactions can be called in an RNA-specific manner. **(c)** *RADIAnT* interaction calling is incorporated into a reads-to-interactions pipeline, for complete processing of diverse RNA-DNA ligation data types. The example output Sankey plot is from a GRID-seq experiment conducted in mouse embryonic stem cells [10]. **(d)** Local *Malat1*-DNA interactions called using *RADIAnT* from GRID-seq [10], RADICL-seq (1% formaldehyde) and RADICL-seq (2% formaldehyde) [8] data generated from mouse embryonic stem cells (mESCs). **(e)** Total numbers of valid paired RNA-DNA reads, unique RNA-DNA interactions and statistically significant (adjusted *p <*0.05) RNA-DNA interactions called by *RADIAnT*, for each replicate of the GRID-seq and RADICL-seq experiments from mESCs. **(f)** Pearson correlation coefficients between adjusted *p* values calculated by *RADIAnT* for common interactions between GRID-seq and RADICL-seq experiments in mESCs, as well as randomly sampled adjusted *p* values, for 5 kb and 25 kb genomic bin sizes.

In a separate approach designed to control for the overrepresentation of specific genomic bins resulting from technical artifacts or sequence biases, *RADIAnT* also constructs a bin-specific background. This is done by calculating the mean RNA interaction frequency at each genomic bin (**Fig. 2a, bottom**). Only interaction frequencies where the RNA in question originates from a different chromosome - and are therefore not subject to linear distance bias - are considered when building bin-specific background.

The two backgrounds are then combined (**Fig. 2a, right**). Consequently, for every genomic bin, there exists a bin-specific background RNA interaction frequency in addition to a distance-based background interaction frequency (imposed if the RNA in question originates from the same chromosome as the bin). Finally, the backgrounds are unified by taking the maximum expected interaction frequency per bin, ensuring maximum stringency.

To call interactions for a given RNA (**Fig. 2b**), the unified background interaction frequency is scaled by the RNA abundance in the input data, giving an expected RNA-bin interaction count. The observed RNA-bin count is compared against the expected RNA-bin count in a Poisson test, resulting in a *p* value for the RNA-bin interaction. The interaction *p* value is adjusted for multiple testing according to the total tested RNA-bin combinations. By default, this encompasses any RNA-bin interaction with at least two supporting reads. Interactions with an adjusted *p* value *<* 0.05 are taken as significant.

*RADIAnT* is incorporated into a reads-to-interactions Snakemake [19] pipeline **(Fig. 2c)**, which takes sequencing reads generated from RNA-DNA ligation experiments as input. The pipeline conducts appropriate pre-processing, and returns called interactions. By implementing the pipeline on different RNA-DNA ligation experiments from the same cell type, the capability of *RADIAnT* to adapt to different input data, whilst maintaining consistent interaction calling, was assessed.

### *RADIAnT* calls consistent RNA-DNA interactions regardless of input data type

To determine the consistency of *RADIAnT*, independent RNA-DNA ligation experiments from the same biological context were utilized. This entailed two triplicate RADICL-seq experiments (1 % and 2 % formaldehyde fixation) [8] and a duplicate GRID-seq experiment [10], all conducted using mouse embryonic stem cells (mESCs).

For an assessment of how consistently *RADIAnT* calls interactions against a predominantly distance-specific background, several megabases surrounding the gene locus of *Malat1* were considered. Malat1 is an extensively studied lncRNA; it is highly conserved, ubiquitously expressed and has important regulatory functions in transcription and alternative splicing [20]. In spite of the different methods, experimental conditions and sequencing depths, a similar pattern of *Malat1*-DNA interactions were called by *RADIAnT* (**Fig. 2d**). Most notably, highly significant interactions were called across a region between 8.75 Mb to 9 Mb in all three datasets.

For a genome-wide perspective, the numbers of valid paired RNA-DNA reads, RNA-bin interactions and statistically significant RNA-DNA interactions called by *RADIAnT* were compared across the experiments. An expected relationship between the number of valid reads and number of significant interactions could be observed (**Fig. 2e**). However, this was not due to increased numbers of interactions in general.

To test genome-wide concordance between called interactions, negative logtransformed adjusted *p* values for common interactions detected (read count *>* 0) across all the experimental replicates were compared. Comparisons were made between RNA-bin interactions at higher (5 kb) and lower (25 kb) resolutions. In each case, the interactions called by *RADIAnT* were strongly correlated across the independent experiments and replicates, except for randomly sampled values which acted as a negative control (**Fig. 2f**). Whilst those interactions called using 25 kb bins were more concordant, those called with 5 kb bins were still consistent across the experiments. Given the likely relationship between sequencing depth and maximal resolution, genomic bin size is a user-defined parameter in the *RADIAnT* pipeline, which can be tuned based on the depth and quality of the data in question.

While *RADIAnT* could call consistent RNA-DNA interactions across different experimental contexts, the question remained whether the interactions called by *RADIAnT* were biologically meaningful, and how well they reflected interactions identified by analogous methods. To approach this, interactions called by *RADIAnT* were compared against those detected in a one-to-all method for detection of RNA-DNA interactions. Additionally, *RADIAnT* interactions were compared to those called by alternative methods proposed for analysis of RNA-DNA ligation data.

### *RADIAnT* accurately detects previously reported *Malat1*-DNA interactions

To assess the ability of *RADIAnT* to identify specific RNA-DNA interactions, RAP-DNA data for *Malat1* in mESCs [17] was utilized. In RAP-DNA, regions of the genome bound by a single RNA, e.g. *Malat1*, can be identified with high accuracy. Peaks called from these data were used to designate whether genomic bins were bound by *Malat1* or not (**Fig. 3a**). *Malat1*-positive and -negative bins were subsequently used to assess the performance of different methods in the detection of *Malat1*-DNA interactions from either RADICL-seq or GRID-seq data generated from mESCs [8, 10]. The methods compared here were *RADIAnT, BaRDIC* [16], the method proposed by Bonetti *et al*. [8] and that proposed by Zhou *et al*. [18]. The final two approaches were reimplemented in R in order to maintain consistency, and the same preprocessing was applied in each case.

**Fig. 3.**
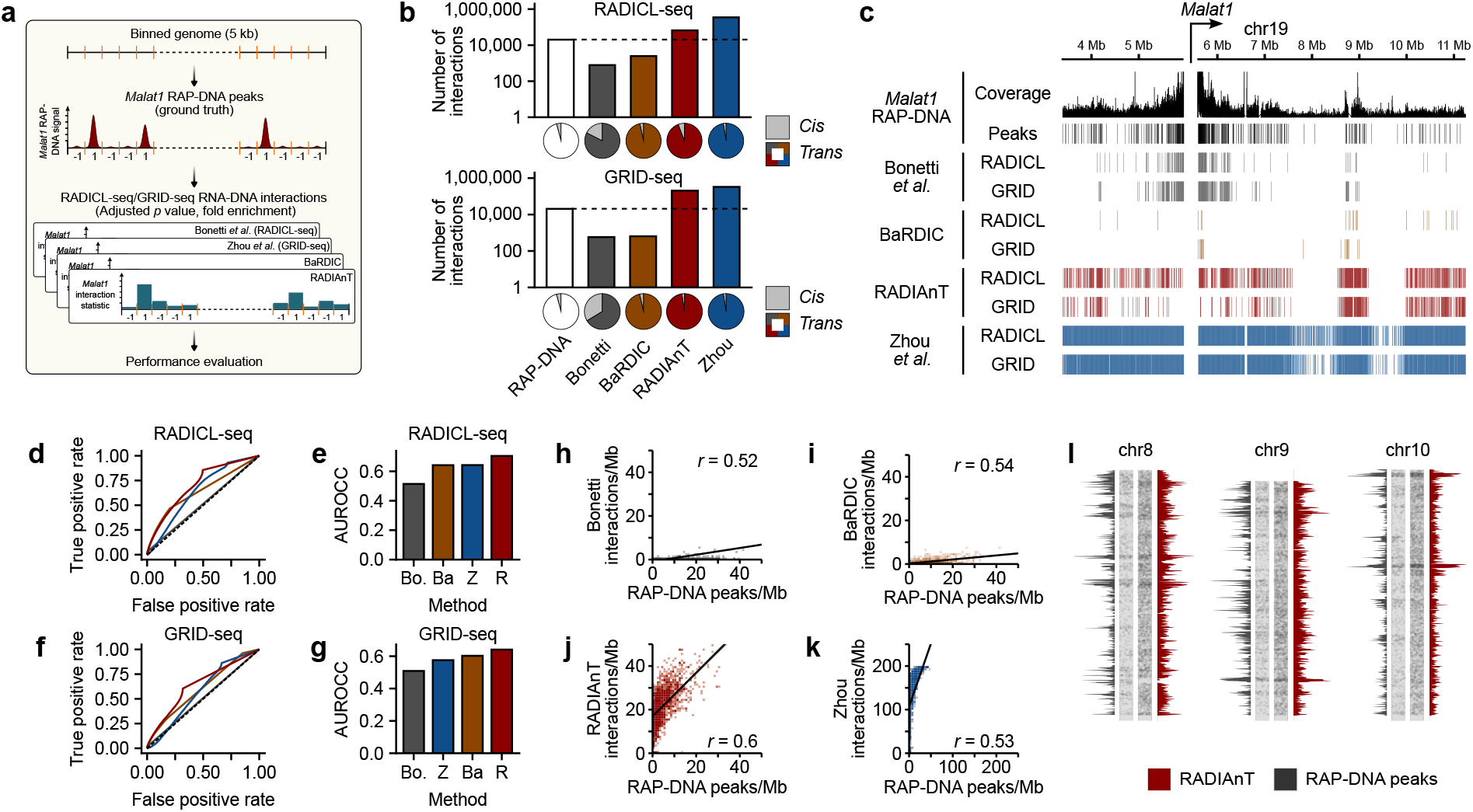
RADIAnT accurately detects *Malat1*-DNA interactions reported in RAP-DNA experiments. **(a)** Workflow for benchmarking of RNA-DNA interaction calling, using *Malat1* RAP-DNA as a ground truth. **(b)** Total *Malat1*-DNA interactions called per method from RADICL-seq and GRID-seq data. **(c)** *Malat1*-DNA interactions called from RADICL-seq and GRID-seq in the vicinity of the *Malat1* gene locus by each method, along with RAP-DNA coverage and peaks. **(d-g)** Receiver operating characteristic analysis per method for the recall of genome-wide RAP-DNA *Malat1*-DNA interactions. **(h-k)** Correlations between total *Malat1*-DNA interactions called per megabase of the genome for each analysis method, compared to total *Malat1* RAP-DNA peaks called per megabase.**(l)** Chromosome scale RADIAnT interaction density compared to *Malat1* RAP-DNA peak density, for chromosomes 8, 9 and 10.

Each method was used to call interactions from both datasets. When comparing total interactions called compared to the number of *Malat1*-positive bins assigned from the RAP-DNA, the Bonetti *et al*. and *BaRDIC* methods called fewer interactions from both RADICL-seq and GRID-seq data (**Fig. 3b**). Conversely, the Zhou *et al*. method called considerably more interactions in both datasets than assigned from the RAP-DNA. *RADIAnT* also called more interactions than were assigned by RAP-DNA, but not to the same extreme. *RADIAnT, BaRDIC* and Zhou *et al*. all detected similar ratios of *cis*-to *trans*-chromosomal *Malat1*-DNA interactions as *Malat1* RAP-DNA, with Bonetti *et al*. detecting a greater proportion of *cis* interactions.

To examine the *cis*-chromosomal interaction in more detail, several megabases surrounding the *Malat1* gene locus were considered (**Fig. 3c**). Here, *BaRDIC* called fewer interactions than would have been expected from the *Malat1* RAP-DNA. In contrast, Zhou *et al*. called interactions across almost the entirety of the region. *RADIAnT* and Bonetti *et al*. called interactions which most closely represented the distribution of *Malat1* RAP-DNA peaks across the region. This indicates that the errant *cis*:*trans* interaction ratio called by Bonetti *et al*. is an outcome of insufficient sensitivity to call *trans*-chromosomal interactions, rather than the calling of bogus *cis* interactions.

To assess the relative performance of each method in a genome-wide manner, a receiver operating characteristic analysis was performed. In both RADICL-seq and GRID-seq, *RADIAnT* outperformed the other methods in the accurate recall of *Malat1*-DNA interactions (**Fig. 3d-g**). Similarly, in a comparison of called interactions per megabase and RAP-DNA peaks per megabase of the genome, *RADIAnT* interactions had a higher correlation coefficient than the other methods (**Fig. 3h-k**). In a visualization of this, *RADIAnT* could identify chromosomal domains enriched in *Malat1* RAP-DNA binding, demonstrating the ability of *RADIAnT* to identify broad RNA-binding regions (**Fig. 3l**).

Having demonstrated that *RADIAnT* can accurately detect genome-wide RNA-DNA interactions, we then turned our attention to its potential for identifying gene regulatory RNAs in a dynamic biological context.

### Detection of dynamic RNA-DNA interactions during EndMT using *RADIAnT*

To test the ability of *RADIAnT* to identify dynamic RNA-DNA interactions between biological settings, endothelial-to-mesenchymal transition (EndMT) was used as a model. In EndMT, endothelial cells transdifferentiate to a more proliferative and invasive mesenchymal-like phenotype, which has been linked to physiological and pathological processes in the cardiovascular system [21, 22]. In an *in vitro* model of EndMT, human umbilical vein endothelial cells (HUVECs) were treated with interleukin-1 beta (IL-1β) and transforming growth factor beta 2 (TGF-β2) to induce a phenotypic switch (**Fig. 4a**). Nuclear RNA-sequencing (nucRNA-seq), RADICL-seq and ATAC-seq were performed on control (endothelial) and treated (EndMT) cells.

**Fig. 4.**
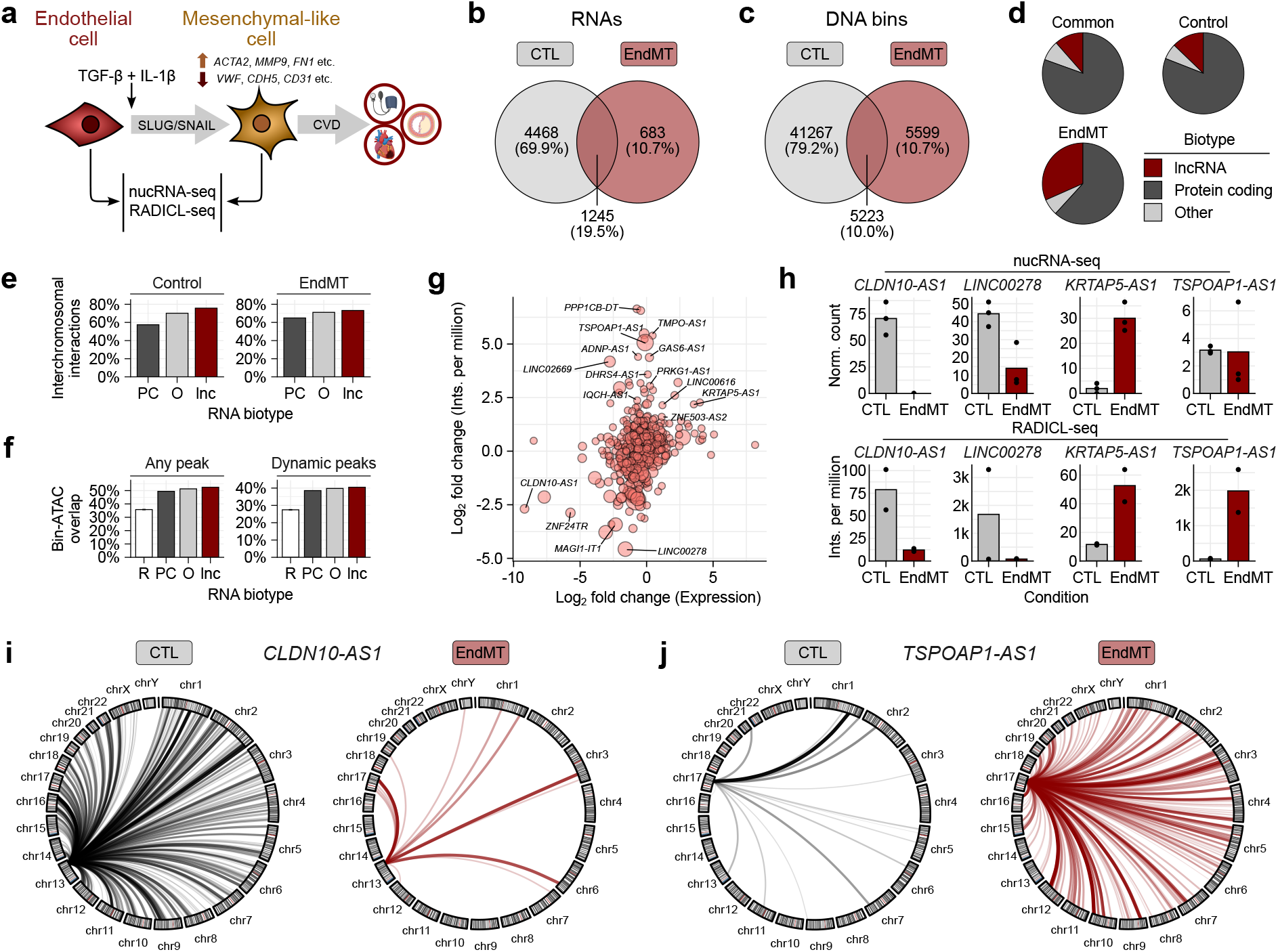
Detection of dynamic RNA-DNA interactions during EndMT using RADIAnT. **(a)** Overview of endothelial-to-mesenchymal transition. **(b-c)** Unique and common interacting RNAs and interacting DNA bins in control and EndMT conditions. **(d)** RNA biotype proportions in control, common and EndMT RNA populations. **(e)** Percentage of interchromosomal RNA-DNA interactions for protein-coding RNAs (PC), long non-coding RNAs (lnc) and other RNA biotypes (O). **(f)** Percentage of RNA biotype-bound DNA bins overlapping with any ATAC-seq peaks detected in control and EndMT conditions, or dynamic ATAC-seq peaks between the conditions. **(g)** LncRNAs stratified by change in expression measured by nucRNA-seq and change in DNA interactions (interactions per million) as detected by RADICL-seq. **(h)** Interaction proportion changes between control and EndMT for the lncRNAs *CLDN10-AS1, LINC00278, KRTAP5-AS1* and *TSPOAP1-AS1*. **(h)** RNA-DNA interactions called for *CLDN10-AS1* in control and EndMT conditions. **(i)** RNA-DNA interactions called for *TSPOAP1-AS1* between control and EndMT conditions.

When comparing RNA and DNA populations involved in *RADIAnT* RNA-DNA interactions detected in two biological replicates of control or EndMT conditions, condition-specific and common features could be extracted **(Fig. 4b-c)**. When examining these populations, there was a greater proportion of lncRNAs involved in interactions in EndMT compared to common or control conditions **(Fig. 4d)**. LncRNAs were also involved in greater proportions of interchromosomal RNA-DNA interactions across control and EndMT condition **(Fig. 4e)**, as well as interacting at a greater proportion of regions where ATAC-seq peaks were detected compared to protein-coding or other RNA biotypes **(Fig. 4f)**.

To identify dynamic DNA-binding lncRNAs in EndMT, nucRNA-seq data were integrated with the RNA-DNA interactions called by *RADIAnT*, with the proportion of the total interactions per condition being used as a metric to measure RNA-DNA binding changes. A number of lncRNAs whose expression and interaction proportion were dynamic between the conditions were detected **(Fig. 4g)**, amongst them *CLDN10-AS1, LINC00278, KRTAP5-AS1* and *TSPOAP1-AS1* **(Fig. 4h)**. When visualized in a genome-wide manner, a dynamic decrease in *CLDN10-AS1*-DNA binding between control and EndMT conditions could be observed **(Fig. 4i)**, compared to an increase in *TSPOAP1-AS1*-DNA binding **(Fig. 4j)**.

This use case suggests that *RADIAnT* can be utilized to examine dynamic biological processes, and identify putative RNA regulators involved in physiological and pathological processes.

## Discussion

Here, we present *RADIAnT*, a statistical method for detection of biologically meaningful interactions from diverse RNA-DNA ligation data types. *RADIAnT* is incorporated into a reads-to-interactions pipeline, and is generalizable across any RNA-DNA ligation method. *RADIAnT* outperformed other proposed methods for calling of analogously-identified RNA-DNA interactions. Tailored to employ any given dataset to build an adequate background, RADIAnT is formulated to be robust to dataset depth and quality. However, it should be recognized that there remain several limiting factors in the generation and analysis of RNA-DNA interactions, which cannot be completely addressed here.

For one, the highly complex state of RNA-DNA interactions coupled with the relative inefficiency of ligation methods in returning valid read pairs (see **Fig. 2c**) results in extremely sparse output data. In the experiments analyzed here, we observed an average efficiency (total reads in called interactions vs. total reads sequenced) of approximately 10%. The largest loss of informative reads is to ribosomal RNA, which often made up half of the total library. Ideally, this would be selected against in the course of the molecular method as in standard RNA-seq [23], however, the biotinylated state of the linker oligonucleotides used in RNA-DNA ligation methods makes this difficult.

The problem of data sparsity could be observed in the benchmarking experiment carried out here. We used *Malat1*-DNA interactions called from RAP-DNA [17], which was sequenced to a depth of approximately 30 million reads. In contrast, the total RADICL-seq or GRID-seq reads involving *Malat1* were approximately 2 million and 1 million, respectively. In the cases of the *BaRDIC* [16] and Bonetti *et al*. [8] methods, this sparsity resulted in severe under-calling of *Malat1*-DNA interactions. Conversely, *RADIAnT* and the Zhou *et al*. [18] method tended to overcall, with *RADIAnT* returning closer numbers of interactions to the RAP-DNA experiment. Given that *Malat1* is amongst the highest expressed genes in the genome, the problem of data sparsity is likely to be magnified when considering more lowly expressed genes. For this reason, we believe that the increased sensitivity displayed by *RADIAnT* is preferable to prioritizing specificity, and subsequently overlooking potentially interesting interactions.

As RNA-DNA ligation methods increase in efficiency and depth, we are confident that the robust and adaptable formulation of *RADIAnT* will continue to be able to effectively detect biologically-relevant RNA-DNA interactions. By building dataset-specific backgrounds, *RADIAnT* will be able to adapt to the decreasing prevalence of confounding factors and artifacts in these data. Additionally, as data sparsity decreases, *RADIAnT* can be tuned to yield higher resolution RNA-DNA interactions. This will permit more accurate integration of RNA-DNA ligation data with other data types and provide greater insight into the biological consequences of RNA-DNA interactions.

In an example use case of *RADIAnT*, we showed that dynamics in RNA-DNA interactions could be detected in the physiologically- and pathologically-important process of endothelial-to-mesenchymal transition. Specific and common interactions were identified per condition. Integration of these with nucRNA-seq permitted the identification of putative gene regulatory lncRNAs which may play a role in defining cellular transitions.

We believe that *RADIAnT* has the potential to become a widely-adopted, standard tool for the unified analysis of a variety of RNA-DNA ligation data types. The formulation of *RADIAnT*, along with our commitment to its maintenance and development, should make it robust to developing molecular methods, and the integration of *RADIAnT* into a Snakemake pipeline minimizes user input and maximizes reproducibility.

## Methods

### Identification of confounding factors

#### Identification of distance bias

RNA and DNA tags of mouse embryonic stem cells sequenced with GRID-seq (GEO accession no.: GSM2396700) and RADICL-seq (GEO accession no.: GSE132192) were mapped to the murine genome assembly version mm39 using *STAR*[24] (v2.7.3a), annotated in GENCODE release M35. RNA and DNA tags of Red-C-sequenced human K562 cells (GEO accession no.: GSE136141) were mapped to the human genome assembly version hg38, annotated in GENCODE release 43. The distance between the tags was computed as the difference between the centre of the 5 kb bin the DNA tag was mapped to and either the start or the end (whichever lies closer) of the gene the RNA tag was mapped to. Gene start/end were rounded to 5 kb. To differentiate reads that captured nascent RNA from other reads, we furthermore differentiated by strand. Cases in which either 1) both tags mapped to the plus strand or 2) both tags mapped to the minus strand (same-strandedness) from cases in which the tags mapped to different strands (opposite-strandedness), assuming that the latter would reflect nascent transcription. Visualisation was produced with ggplot2 (v3.5.0).

#### Analysis of RNA abundance

To compare interaction frequency as captured by RADICL-seq, GRID-seq and Red-C against gene expression, we mapped sequencing reads from CAGE (GEO accession no. GSE132191), GRO-sequencing (GEO accession no. GSE82312) and RNA-sequencing (GEO accession no. GSE136141), respectively, to the genomes as described in the section above using *STAR*[24] (v2.7.3a) and quantified using *featureCounts* (v2.0.0). Statistical analysis was conducted in R (v4.3.1), visualisation with ggplot2 (v3.5.0).

### Preprocessing

#### Depletion of ribosomal RNA

Potential contaminating ribosomal RNA is eliminated with *BBDuk* of the *BBTools* suite [25]. In our analysis of the data presented in this work, we used *BBTools* (v39.01) and the non-redundant SSU and LSU Ref datasets of the SILVA [26] (release 138.1) as a ribosomal reference database.

#### Mapping

The split RNA and DNA tags (FASTQ format) are aligned to the genome using *STAR*[24] (v2.7.3a). Apart from default parameters, we set `--alignIntronMax 1` to disable splicing, `--outFilterScoreMinOverLread 0` and `--outFilterMatchNminOverLread 0` and `--outFilterMatchNmin 0` to enhance the mapping rate. Multi-mapping reads are discarded using *SAMtools* [27] (v1.10).

#### Identification of interacting regions

To describe interactions, the genome is split into two different kinds of units: For the chromatin part of an interaction, we regard bins of size 5 kb as the smallest unit of the genome. For the RNA part of an interaction, we regard genes as the smallest unit of the genome. *BEDtools* [28] *intersect* (v2.27.1) is used to assign DNA tags to bins and RNA tags to genes, respectively. Since multiple genes can be annotated to the same locus, the read is assigned to the gene it has the maximum overlap with.

#### Distance Computation

For intrachromosomal interactions, an interaction distance can be computed. Annotated gene starts and ends are rounded to match the center of the closest 5 kb bin. The distance is computed between either the rounded end or rounded start of the gene (whichever lies closer), and the center of the DNA bin it interacts with.

### Statistics

#### Background model construction

To account for experimental artifacts, we build a distance- and expression-sensitive background to model the expected interaction frequency of a given bin with a given RNA. The distance-specific background is calculated for each RNA *r* as the number of reads that were sequenced at a given distance, normalized by the total number of reads for RNA *r*. For each given distance, the interaction frequency is given by the mean over all RNAs:

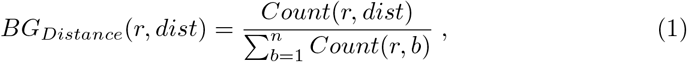

where *n* is the total number of bins on the chromosome where RNA *r* resides. *Count*(*r, dist*) denotes the number of RNA-DNA interaction counts that RNA *r* has 13 with the bin *b* at distance *dist* to the gene locus of *r. Count*(*r, b*) denotes the number of RNA-DNA interaction counts that RNA *r* has with the bin *b*.

Since there are cases in which no distance can be computed (e.g. cross-chromosome interactions), a whole-genome bin-specific background is constructed in the same manner:

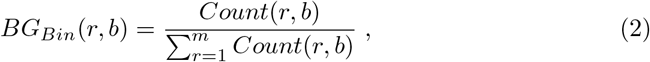

where *m* is the number of all bins on all chromosomes. Regions which are not covered by reads of an RNA *r* are by definition set to *Count*(*r, b*) = 0.

The two background models are unified in a manner that favors the more conservative value for later testing:

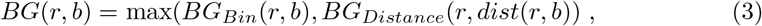

where *dist*(*r, b*) denotes the distance of the bin *b* to the gene locus of *r*.

#### Identification of statistically robust RNA-DNA interactions

A particular RNA-DNA interaction *count*(*r, b*) is assessed for statistical significance. Here an one-tailed Poisson test is used. The mean *λ* is defined by incorporating the background count and the estimated RNA abundance value:

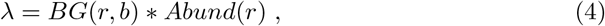

where *Abund*(*r*) denotes the estimated RNA abundance in the dataset by summing over all RNA-DNA interactions of *r*:

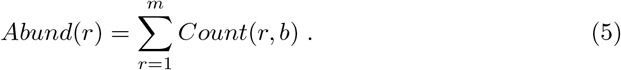

We conduct a Poisson test using significance level of *α* = 0.05. To account for multiple testing, we apply the method described by Benjamini & Hochberg (1995)[29] and compute the corresponding FDR.

#### Snakemake pipeline assembly

*RADIAnT* is made publicly available as a Snakemake [19] pipeline. Input to the pipeline are split RNA and DNA FASTQ files, and a configuration file which can be modified by the user. The pipeline executes all necessary pre-processing steps (depletion of ribosomal RNA, elimination of multimapping reads, validation of complete RNA-DNA pairs). If Red-C has been specified as the sequencing method, *RADIAnT* accounts for separate processing of the 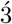 and 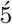 parts of the RNA read in the aforementioned steps. The Snakemake pipeline subsequently applies the *RADIAnT* statistical method on the resulting set of valid RNA-DNA pairs. The outputs of the pipeline are RNA-DNA interactions called by *RADIAnT*, as well as several plots detailing important metrics and quality checks for the dataset.

### Assessing the consistency of *RADIAnT* interactions

#### Datasets

For the comparison of RNA-DNA interaction called by *RADIAnT* across different RNA-DNA ligation methods and experiments, RADICL-seq [8] and GRID-seq [10] data generated in mouse embryonic stem cells (mESCs) were downloaded and analyzed. The NCBI GEO accession numbers for the RADICL-seq and GRID-seq data are GSE132192 and GSE82312, respectively, with there being two independent RADICL-seq experiments, one with 1 % formaldehyde fixation and one with 2 %.

#### Interaction calling with *RADIAnT*

Each of the independent replicates of each experiment was input to the *RADIAnT* Snakemake pipeline, and analyzed with annotation data for the mm39 (GENCODE M35) genome assembly and 5 kb genomic bins. Interactions were then compared against one another locally by considering *Malat1*-DNA interactions in the genomic region surrounding the *Malat1* gene locus. Background interaction frequency, *Malat1* interaction frequency, and sites of significant (adjusted *p <* 0.05) *Malat1*-DNA interactions were plotted using ggplot2 per RNA-DNA ligation experiment.

#### Correlation between *RADIAnT* interactions per RNA-DNA ligation method

To test the consistency of interactions called by *RADIAnT* between RNA-DNA ligation methods, common interactions detected with at least 1 supporting read in each experimental replicate were determined. The adjusted *p* values calculated for these interactions by *RADIAnT* in each experimental replicate was then negative log-transformed, and the resulting values used as input to the R function *cor*.*test* in a pairwise manner. As a negative control, log-transformed adjusted *p* values were sampled in triplicate from the distribution of values calculated across the common interactions for all experimental replicates and assigned to the common interactions as a negative control. This process was carried out for *RADIAnT* interactions called against 5 kb and 25 kb bin sizes. Resulting pairwise Pearson correlation coefficients were plotted using ggplot2 (v3.5.0).

### Benchmarking of RNA-DNA interaction calling

#### *Malat1* RAP-DNA data analysis

Raw data corresponding to *Malat1* RAP-DNA and input data generated from mESCs from the NCBI GEO accession GSE55914 were downloaded and re-processed. Reads were aligned to the mm39 (GENCODE release M35) genome assembly using Bowtie2 [30] (v2.4.5) with the parameters `--local` and `--very-sensitive`. Coverage tracks were generated using bamCoverage [31] (v3.5.3) with the parameter `--normalizeUsing CPM`. Peaks were called using MACS [32] (v3.0.0), with the *Malat1*-RAP alignments as treatment and the input as control.

#### RNA-DNA ligation data

For the comparison between RNA-DNA interaction calling methods, RADICL-seq [8] and GRID-seq [10] data generated in mESCs were downloaded and analyzed with each method. The NCBI GEO accession numbers for the RADICL-seq and GRID-seq data are GSE132192 and GSE82312, respectively.

#### Analysis with *RADIAnT*

Both RADICL-seq and GRID-seq reads were used as input to the *RADIAnT* Snake-make pipeline in order to call interactions. In both cases, the mm39 (GENCODE M35 release) genome assembly was used in combination with 5 kb genomic bins. RNA-bin interaction counts were pooled across replicates prior to interaction calling with *RADIAnT*. Statistically significant interactions were designated as those with an adjusted *p <* 0.05.

#### Implementation of Bonetti et al. statistical method

The computational method for calling RNA-DNA interactions as proposed by Bonetti *et al*. [8] was reimplemented in R. Briefly, RNA-bin interactions were subjected to a binomial test where the parameter were as follows: *x* was the RNA-bin interaction count, *n* was the sum of all RNA-bin interaction counts for the RNA in question, *p* is the reciprocal of the total number of bins detected as being bound by the RNA, and *alternative* was set to “greater”. Resulting *p* values were adjusted for multiple testing using the Benjamini-Hochberg procedure [29].

#### Implementation of Zhou et al. statistical method

The computational method for calling RNA-DNA interactions as proposed by Zhou *et al*. [18] was also reimplemented in R. Briefly, RNA-bin interactions were subset to *trans*-chromosomal interactions of mRNAs only. A bin-specific background is constructed from the number of reads at each 1kb bin, which is normalized by the overall number of genes in the interaction set. The normalized read count is then smoothed in a moving window approach (10 bins), further normalizing by total read count per chromosome and number of bins per chromosome, yielding a bin-specific background coverage. The approach by Zhou et al. calculates the fold enrichment as the ratio of the observed RNA-bin interaction frequency (normalized by read count per chromosome and number of bins per chromosome) to the background coverage.

#### Calling of RNA-DNA interactions using *BaRDIC*

*BaRDIC* was installed and run as per the instructions at https://github.com/dmitrymyl/BaRDIC/. Genomic annotation and chromosome sizes were supplied based on mm39/GENCODE M35, and the background RNAs used were the 100 most abundant mRNAs present in the dataset.

#### Performance evaluation

Each RNA-DNA interaction calling method was implemented as described above. RADICL-seq and GRID-seq data were subjected to identical pre-processing procedures prior to interactions being called by each method. Performance evaluation of each of the tools was carried out in R using the ROCR (v1.0.11) package. The evaluation dataset was composed of 5 kb genomic bins which either intersected with a *Malat1*

RAP-DNA peak or not. For each method, a predictor value was assigned per bin. These values were negative log-transformed adjusted *p* values for *RADIAnT, BaRDIC* and Bonetti *et al*., and fold enrichment for Zhou *et al*.. The relative performance of each tool was then computed with ROCR, and receiver operating characteristic (ROC) curves were plotted with ggplot2, along with area under the ROC curve.

Additionally, genome-wide concordance between interactions called per method and *Malat1* RAP-DNA was computed. This was done by taking comparing the total interactions called per megabase total *Malat1* RAP-DNA peaks per megabase. Similarly, *RADIAnT* interaction density was compared to *Malat1* RAP-DNA peak density on a chromosomal scale, and was plotted with ggplot2.

### Cell culture

Human umbilical vein endothelial cells (HUVEC, batches 405Z013, 408Z014, 416Z042, Lonza) were cultured on 0.2 % gelatin-coated dishes in endothelial growth medium (EGM, Pelo Biotech). EGM was supplemented with 8 % fetal calf serum (FCS, Sigma-Aldrich), 50 U*/*mL penicillin (Gibco) and 50 µg*/*mL streptomycin (Gibco). The cells were thawed freshly and passaged once on a 10 cm dish (Sarstedt). Experiments were performed with passage 3. Cells were incubated at 37 ^*°*^C with 5 % CO_2_. If not stated otherwise, three batches of HUVECs were used for the experiments. All media used were pre-warmed at 37 ^*°*^C.

### Endothelial-to-mesenchymal transition

Endothelial-to-mesenchymal transition (EndMT) was induced according to [33]. HUVECs were seeded in EGM (Pelo Biotech) with a density of 6900 cells*/*cm^2^ on 0.2 % gelatin-coated dishes (15 cm, Sarstedt) one day prior to induction of EndMT. To induce EndMT, differentiation medium (DM) was used, which consisted of endothelial basal medium (EBM, Pelo Biotech) supplemented with 8 % FCS (Sigma-Aldrich), 50 U*/*mL penicillin (Gibco), 50 µg*/*mL streptomycin (Gibco) and L-glutamine (Gibco). 10 ng*/*mL TGF-β2 (Peprotech) and 1 ng*/*mL IL-1β (Peprotech) were added freshly to pre-warmed DM medium. Medium was changed every 24 h for a total of 5 days.

### Nuclear RNA sequencing

Nuclear RNA sequencing was performed on control HUVECs and HUVECs stimulated to undergo EndMT. To isolate nuclear extract, medium was aspirated and cells were washed once with cold Hank’s solution. Harvesting was performed by scraping cells twice in 500 µL Hank’s solution and centrifuging (1 min, 13 000 rpm at 4 ^*°*^C). The supernatant was removed and the pellet resuspended in 600 µL buffer A (10 mM HEPES pH 7.9, 10 mM KCl, 0.1 mM EDTA, 0.2 mM EGTA, 1 mM DTT, 40 µg*/*mL PMSF, 12 µg*/*mL protease inhibitor mix). After incubation on ice for 15 minutes, 45 µL 10 % Nonidet (Sigma Aldrich) were added to lyse the cytosolic membrane, the cells were vortexed for 10 seconds and centrifuged (1 min, 13 000 rpm at 4 ^*°*^C). The supernatant, containing the cytosolic fraction, was removed and the pellet resuspended in 150 µL buffer C (20 mM HEPES pH 7.9, 0.4 M NaCl, 1 mM EDTA, 1 mM EGTA, 1 mM DTT, 40 µg*/*mL PMSF, 12 µL*/*mL protease inhibitor mix). To lyse the nuclei, the resuspended pellet was rocked for 15 minutes at 1400 rpm and 4 ^*°*^C. To remove insoluble debris, the lysate was centrifuged (5 min, 13 000 rpm at 4 ^*°*^C) and the supernatant was transferred to a new 1.5 mL tube.

RNA isolation was performed according to Bio&SELL RNA isolation kit manual. If not otherwise stated, RNA was eluted in 30 µL at 13 000 rpm for 1 minute at room temperature. RNA concentration was measured with the NanoDrop ND-1000. Library preparation was carried out using the SMARTer^®^ Stranded Total RNA-Seq Kit v2 – Pico input mammalian. Libraries were sequenced on an Illumina NextSeq 500 sequencer with single-end, 75 bp read length and to an average depth of 30 million reads.

### Nuclear RNA sequencing analysis

Nuclear RNA sequencing reads were aligned to the hg38 (GENCODE v43) genome assembly using STAR [24] (v2.7.10) with the parameters `--outSAMtype BAM SortedByCoordinate --quantMode GeneCounts`. Differential gene expression analysis was performed using DESeq2 (v1.40.2) [34], with the respective donor per sample included in the experimental design.

### Assay for transposase accessibility (ATAC)-sequencing

ATAC resuspension buffer was prepared (10 mM Tris-HCl pH 7.4, 10 mM NaCl, 3 mM MgCl_2_). 50 000 cells per biological replicate for control and EndMT conditions were washed once with 50 µL ice-cold PBS and lysed in 50 µL cold ATAC lysis buffer (97 % ATAC resuspension buffer, 0.1 % NP-40, 0.1 % Tween20, 0.01 % Digitonin). Cells were incubated for 3 minutes on ice, before being topped up to 1 mL with ATAC wash buffer (99 % ATAC resuspension buffer, 0.1 % Tween20) and inverted three times. Nuclei were pelleted by centrifugation (10 minutes, 500 *× g*, 4 ^*°*^C). Supernatants were removed and nuclei were resuspended in 50 µL ATAC transposition reaction mix (25 µL 2x Tagment DNA buffer (Illumina), 2.5 µL Transposase (Illumina), 22.5 µL nuclease-free water). After incubation (37 ^*°*^C, 1000 rpm, 30 minutes), the DNA was purified using the Qiagen MinElute kit and eluted in 10 µL Elution buffer. The library was prepared with the ATAC library amplification reaction mix, as described in [35]. Following amplification, the library was sequenced on an Illumina NextSeq 500 sequencer with paired-end, 75 bp read length and to an average depth of 15 million reads.

Resulting sequencing reads were aligned to the hg38 genome assembly (GENCODE v43) using Bowtie2 [30] (v2.4.5) with the parameters `--local` and `--very-sensitive`. Coverage tracks were generated using bamCoverage [31] (v3.5.3) with the parameter `--normalizeUsing CPM`. Dynamic ATAC-seq peaks between control and EndMT conditions were detected using MACS[32] (v3.0.0) `bdgdiff`, with filtered sequencing depths per condition used to normalize between samples.

### RADICL-seq

RADICL-seq was performed as reported in [8] on control HUVECs and HUVECs subjected to EndMT. For each condition, 4 million cells were washed once with PBS, trypsinated and spun down (1200 rpm, 4 minutes). The cell pellet was resuspended in 5 mL PBS and the cells were counted using a Neubauer counting chamber and Trypan blue staining. 2 million cells were cross-linked through resuspension in 2 mL fresh 1 % formaldehyde for 10 minutes at room temperature. Crosslinking was stopped by adding 715 µL of freshly prepared 1 mol*/*L glycine in double-distilled water. Cells were centrifuged (100 *× g*, 5 minutes, 4 ^*°*^C), and the cell pellet was washed once with 5 mL ice-cold PBS and spun down again (100 *× g*, 5 minutes, 4 ^*°*^C). Pellets were shock-frozen in liquid nitrogen and stored at *™* 80 ^*°*^C. Thereon, the RADICL-seq protocol was carried out as described by Bonetti *et al*. [8]. Final library concentrations were measured by Qubit fluorometer and library quality and size was determined by a Bioanalyzer High Sensitivity DNA Analysis according to manufacturer’s protocol (Agilent). Paired-end sequencing with 150 bp read length was carried out using an Illumina NextSeq 500 sequencer with a sequencing depth of 125 million reads per condition.

### Analysis of EndMT RADICL-seq data

Control and EndMT RADICL-seq data were analysed using the *RADIAnT* Snake-make pipeline, using the hg38 genome assembly and 5 kb genomic bins. Significant RNA-DNA interactions were those with a *RADIAnT p* value *<* 0.05. RNA biotypes were assigned to populations of interacting RNAs as annotated in GENCODE release 43.Interaction proportions per RNA were calculated per condition by taking the number of significant interactions per RNA divided by the total number of significant interactions in the condition. Circular genome plots were made using ggplot2 (v3.5.0).

## Supplementary information

## Acknowledgements

We would to extend our gratitude to Dr. Stefan Günther (Max Planck Institute for Heart and Lung Research, Bad Nauheim, Germany) and his group, as well as Professor Bernhard Brüne (Goethe University Frankfurt, Institute of Biochemistry I) and his group, for allowing access to their next-generation sequencing facilities.

## Declarations

### Data availability

Data generated for this manuscript are available at NCBI GEO under the accessions GSE273203 (RADICL-seq), GSE273205 (ATAC-seq) and GSE273207 (nucRNA-seq).

### Code availability

The code for the method described herein is available at https://github.com/si-ze/RADIAnT

### Author contribution

S.Z., M.H.S. and T.W. conceived the idea for the study. S.S. and A.B. planned and performed the wet-lab experiments. R.P.B., M.S.L. and S.S. provided key data and input. S.Z. and T.W. analyzed the data. S.Z. and T.W. wrote the code and formulated the manuscript. All listed authors read, commented on and edited the manuscript.

